## Supplementary Table 1 for "Structural landscape of the Respiratory Syncytial Virus nucleocapsids"

**Supplementary Table 1. Primer sequences used to generate pN deletion and mutations**

| Plasmid name | Primer sequence (5'-3') |
|---|---|
| pN1-370 | Forward ACTAGACTTGtaaGCAGAAGAACTAG<br>Reverse ACACTGTAGTTAATCACAC |
| pNH100E | Forward TGTAACAACAgagCGTCAAGACAT<br>Reverse CTACTCCATTTGCTTTTAC |
| pNH100E-R101D | Forward AACAACAGAGgacCAAGACAT<br>Reverse ACATCTACTCCATTTGCTTTTAC |
| pNH100E-E122R | Forward CTTAACAACTgacATTCAAATCAAC<br>Reverse CTTGCCAATGTTAACACTTC |

The pNH100E plasmid was used as a template to generate the plasmids pNH100E-R101D and pNH100E-E122R.
