## Supplementary Table 2 for "Structural landscape of the Respiratory Syncytial Virus nucleocapsids"

### Supplementary Table 2. Cryo-EM data collection, refinement and validation statistics

| Dataset | WT RSV NC | | | | | Mutant 1-370 RSV NC | |
|---|---|---|---|---|---|---|---|
| **Name of the maps generated in this study** | **N10 double ring** | **Helical NC** | **Ring-capped NC** | **Double-headed NC** | **Helical sub-section** | **Helical NC** | **Stack** |
| **EMDB**<br>**PDB** | EMDB<br>PDB | EMDB | EMDB | EMDB | EMDB<br>PDB | EMDB | EMDB<br>PDB |
| **Data collection and processing** | | | | | | | |
| Magnification | 36,000 | | | | | 36,000 | |
| Voltage (kV) | 200 | | | | | 200 | |
| Electron exposure (e–/Å$^2$) | 42.0 | | | | | 42.0 | |
| Defocus range (μm) | -0.7 – -2.4 | | | | | -0.7 – -2.4 | |
| Pixel size (Å) | 1.145 | | | | | 1.145 | |
| Symmetry imposed | D10 | Helical<br>149.5°<br>105.3Å | C1 | D1 | C1 | Helical<br>-36°<br>6.58Å | D10 |
| Initial particle images (no.) | - | 1406835 | - | - | - | 471,549 | - |
| Final particle images (no.) | 47,212 | 389,540 | 22,162 | 25,338 | 389,540 | 329,706 | 81,918 |
| Map resolution (Å)<br>FSC threshold | 2.9<br>0.143 | 6.2<br>0.143 | 3.9<br>0.143 | 3.8<br>0.143 | 3.5<br>0.143 | 4.3<br>0.143 | 2.8<br>0.143 |
| Map resolution range (Å) | 2.5-3.7 | 5-9 | 3-11 | 3.5-9.5 | 2.5-8.5 | 3.8-7.8 | 2.4-4.4 |
| **Refinement** | | | | | | | |
| Initial model used (PDB code) | 2WJ8 | - | - | - | 2WJ8 | - | 2WJ8 |
| Model resolution (Å) (masked)<br>FSC threshold | 2.9<br>0.5 | - | - | - | 3.7<br>0.5 | - | 2.98<br>0.5 |
| Map sharpening *B* factor (Å$^2$) | -96 Å$^{-2}$ | - | - | - | -94 Å$^{-2}$ | - | -99 Å$^{-2}$ |
| Model composition<br>Non-hydrogen atoms<br>Protein residues<br>Ligands | 61,800<br>7,560<br>0 | - | - | - | 15,415<br>1890<br>0 | - | 120,560<br>14,760<br>0 |
| *B* factors (Å$^{-2}$)<br>Protein | 38.96 | - | - | - | 54.50 | - | 47.85 |
| R.m.s. deviations<br>Bond lengths (Å)<br>Bond angles (°) | 0.005<br>0.628 | - | - | - | 0.007<br>1.021 | - | 0.005<br>0.883 |
| Validation<br>MolProbity score<br>Clashscore<br>Poor rotamers (%) | 1.50<br>8.30<br>0.00 | - | - | - | 2.99<br>18.34<br>15.87 | - | 2.18<br>8.64<br>5.19 |
| Ramachandran plot<br>Favoured (%)<br>Allowed (%)<br>Outliers (%) | 97.77<br>2.23<br>0.00 | - | - | - | 95.48<br>4.52<br>0.00 | - | 97.00<br>3.00<br>0.00 |
