## Supplementary Figures for "Structural landscape of the Respiratory Syncytial Virus nucleocapsids"

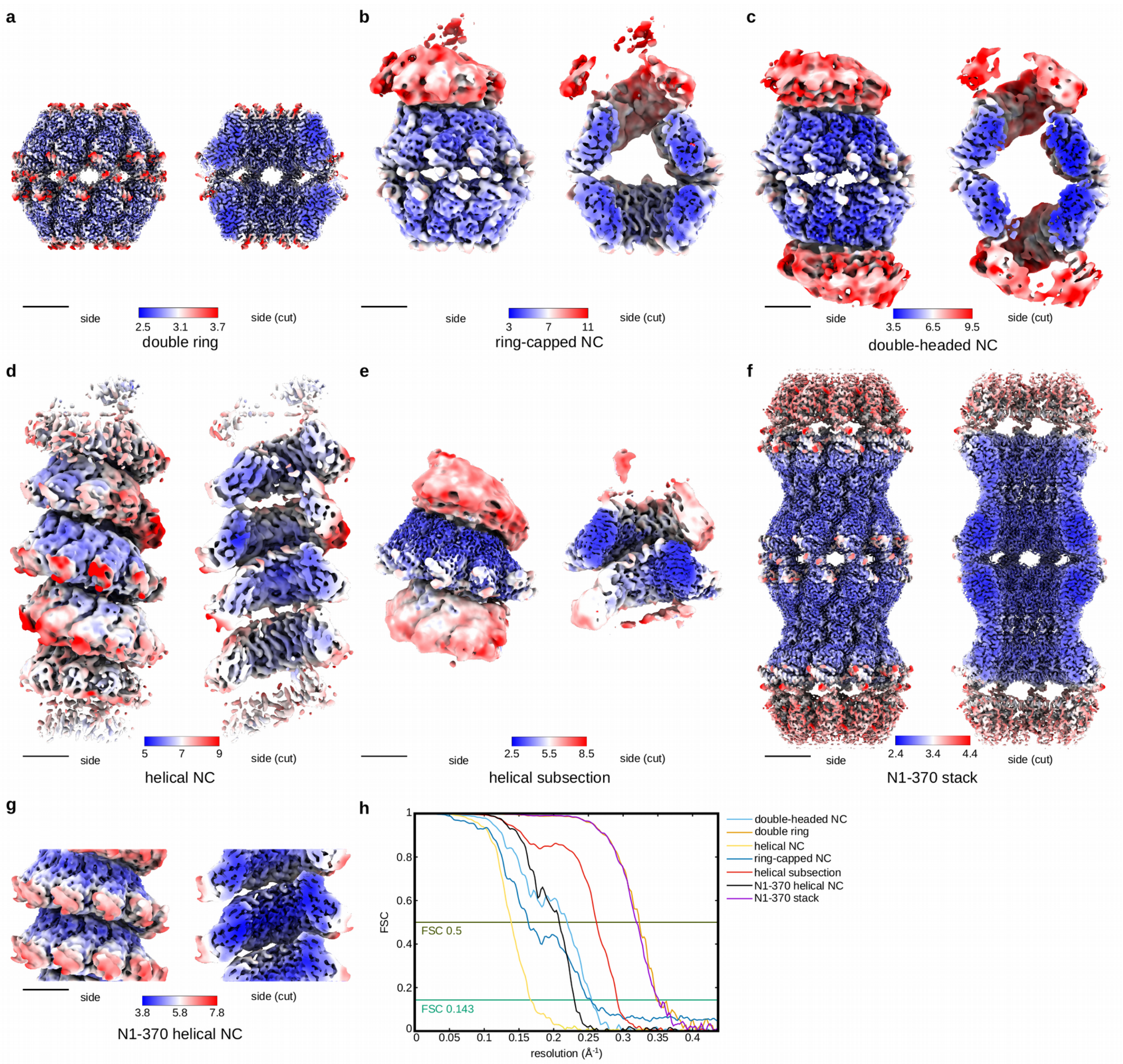


**Supplementary Figure 1. Cryo-EM maps determined in this work and the corresponding FSC curves.** Cryo-EM maps filtered and coloured by local resolution (in Å) for the double ring (a), ring-capped NC (b), double-headed NC (c), helical NC (d), helical subsection (e), N1-370 stack (f) and N1-370 helical NC (g). Contours levels used in ChimeraX to generate the full or cut-out views are 0.264 (a), 0.096 (b), 0.103 (c), 0.117 (d), 0.084 (e), 0.174 (f) and 0.075 (g). Scale bars 50 Å. (h) Masked FSC curves. Scale bars 50 Å.

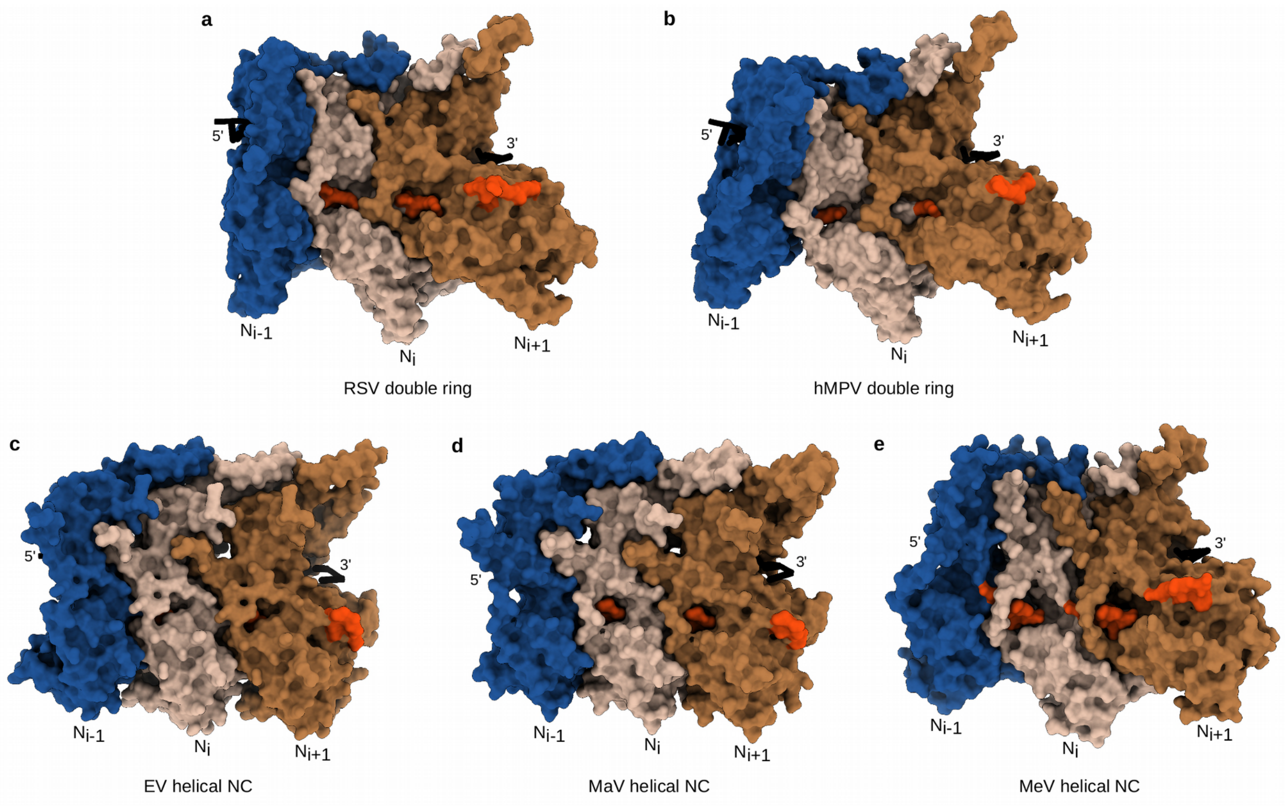


**Supplementary Figure 2. N-hole conservation between *Pneumoviridae*, *Filoviridae* and *Paramyxoviridae* families.** Atomic models of three consecutive promoters of the cryo-EM structure of RSV double ring (a), crystal structure of the hMPV double ring (PDB: 5FVC) (b), Ebola virus helical NC (PDB: 5Z9W) (c), Marburg virus helical NC (PDB: 7F1M) (d), and Measles virus helical NC (PDB: 4UFT) (e) are shown as a surface. The residues coloured in orange correspond to the short loop from the NTD of the $N_{i-1}$ protruding into the N-hole of $N_i$, numbered 230-238 in RSV N (a), 234-238 in hMPV N (b), 221-226 in EV N (c), 203-208 in MaV N (d) and 242-248 in MeV (e). Of note, EV N-hole is less visible because the short loop (in orange) protruding into the N-hole is masked by the loops forming the N-hole. RNA is in black.

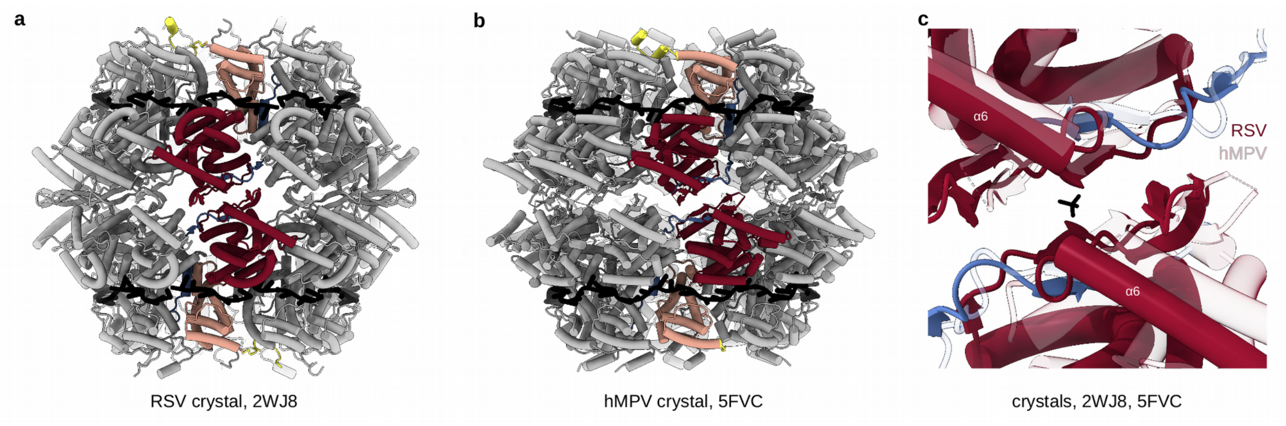


**Supplementary Figure 3. Comparison of RSV and hMPV double rings as observed in the respective crystals.** (a) Atomic model of the RSV $N_{10}$ double ring (PDB: 2WJ8) is shown as cartoon, coloured as in Figure 1. (b) Atomic model of the hMPV $N_{10}$ double ring (PDB: 5FVC). (c) Close-up of the alignment of the RSV and hMPV $N_{10}$ double rings. RSV $N_{10}$ double ring is coloured as in (a) and hMPV $N_{10}$ double ring is in transparent. The borate ion in the interaction site in PDB: 2WJ8 is shown. The $6^{th}$ alpha-helix is labeled for reference.

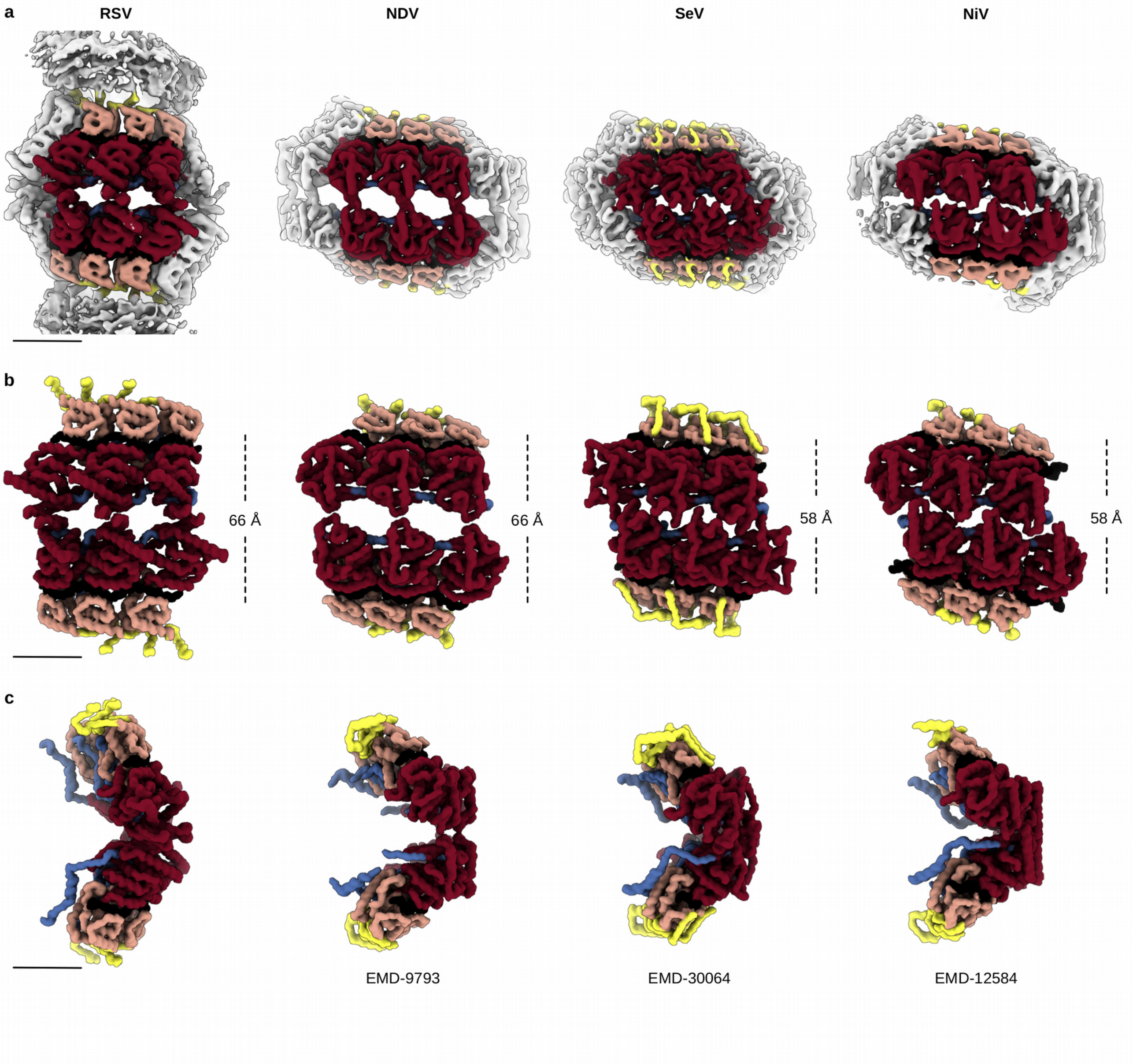

**Supplementary Figure 4. Comparison of double-headed NCs of RSV, NDV, SeV and NiV.** (a) Cryo-EM maps of double-headed NCs of RSV, New-castle disease virus (NDV) (EMD-9793), Sendai virus (SeV) (EMD-30064) and Nipah virus (NiV) (EMD-12584), filtered to 6 Å resolution, scale bar 50 Å. Front (b) and side (c) views of atomic models of three consecutive protomers from the double-headed NCs of RSV, NDV, SeV and NiV, filtered to 6 Å resolution and displayed as surface. Of note, the clam-shaped assemblies of RSV and NDV are both formed in an end-on fashion, without insertion into the opposite inter-protomer grooves and with the centers of gravity of the two spirals separated by 66 Å, as in the cryo-EM structure of the RSV $N_{10}$ double ring. In contrast, the SeV and NiV clams adopt a closely nested packing, leading to inter-spiral distances of only 58 and 56 Å respectively, and therefore actually look more reminiscent of the crystal structure of the RSV double ring. A visual comparison of the side views of the central clam-forming protomers, opposite to the helical junction, also reveals that the the RSV protomer is notably less tilted from the filament axis than its three paramyxoviral counterparts.

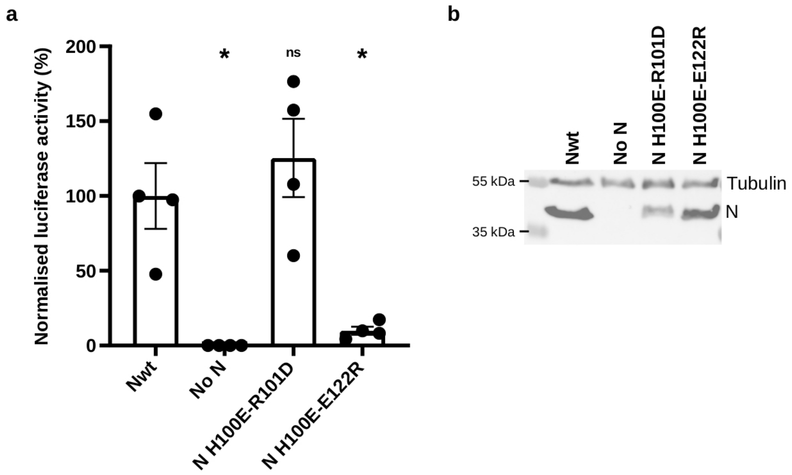


**Supplementary Figure 5. Impact of N mutation on the RSV polymerase activity**. (a) Quantification of the RSV polymerase activity in the presence of wild type N (Nwt), in the absence of N (No N) and for the two N mutants, H100E-R101D and H100E-E122R. Luciferase activities were normalised based on corresponding ß-Gal activities, and are representative of three experiments performed in quadruplicates. Data are mean +/- s.e.m. ( standard error of the mean). n = 4 replicates. A two-tailed Mann-Whitney test was used to compare the normalised luciferase activities between the Nwt and the other conditions. $p < 0.05$ is considered significant. *, p = 0.0286. For N H100E-R101D, p = 0.3429 (ns for no significance). (b) Western blot showing the expression of N (WT and mutants) and tubulin in BSRT7/5 cells.

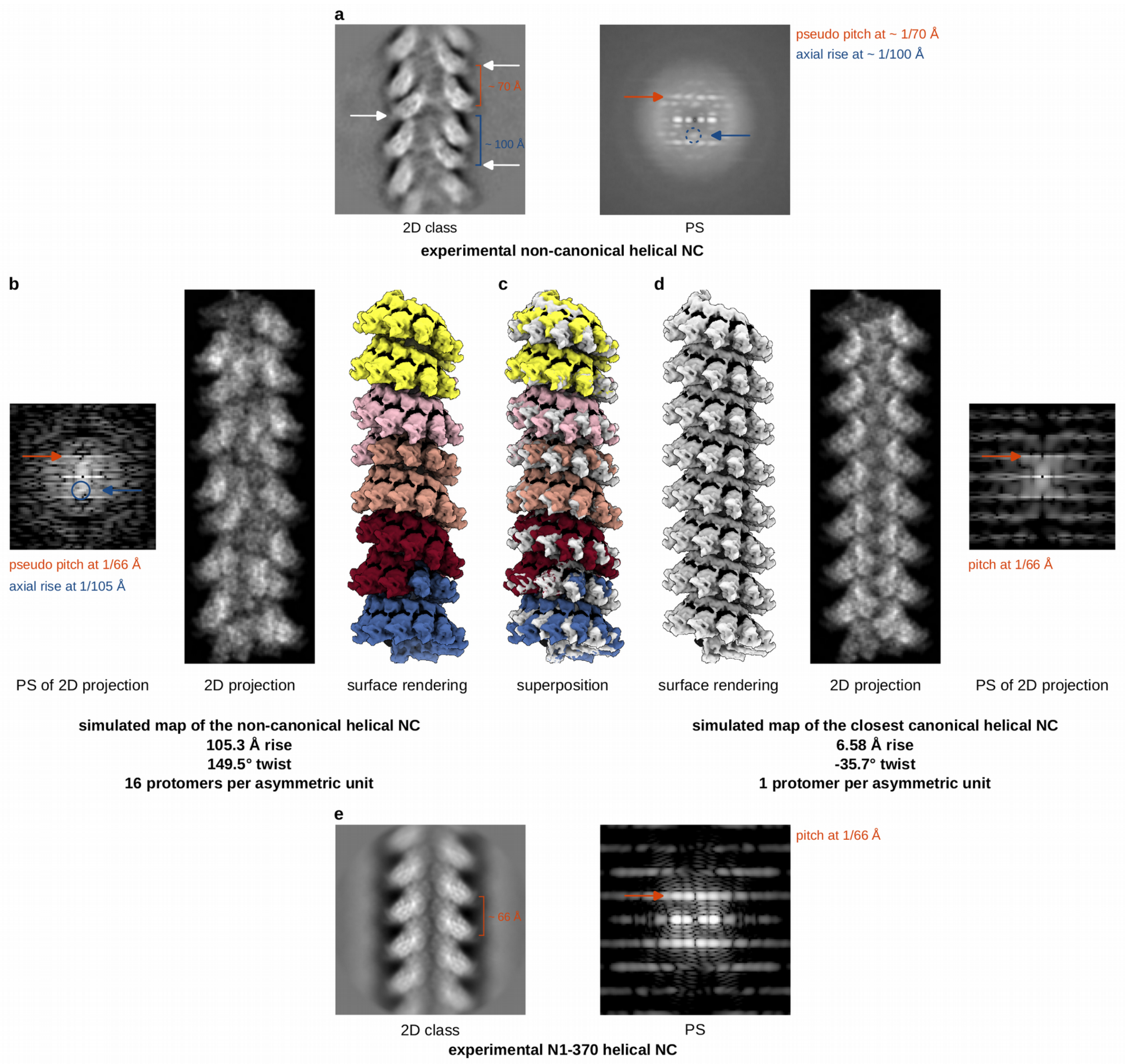


**Supplementary Figure 6. Comparison between the experimental NC, the simulated non-canonical helical NC, the simulated closest canonical helical NC and the N1-370 mutant NC**. (a) A 2D class-average (left) and an average of the PS of the particles in the class (right), representative of the experimental non-canonical helical NCs are shown. The white arrows are indicating the inwards-shifted protomers. (b) A simulated non-canonical helical NC map has been built using the experimentally refined helical parameters as indicated and filtered to 8 Å resolution (each asymmetric unit is coloured in a different colour, right). The corresponding 2D projection (middle) and its PS (left) are shown. (c) Superposition of the simulated maps of the non-canonical helical NC and the closest canonical helical NC. (d) The closest canonical helical NC symmetry parameters were calculated by dividing the non-canonical helical rise and twist by the number of protomers in the asymmetric unit (16; 105.3/16=6.58 Å rise and a -(360-149.5+360)/16=-35.7° twist to account for right-handed to left-handed conversion). The corresponding simulated map filtered to 8 Å resolution (left), its 2D projection (middle) and its PS (right) are shown. (e) A 2D class-average (left) and an average of the PS of the particles in the class (right), representative of the canonical helical NC formed by the N1-370 mutant construct.

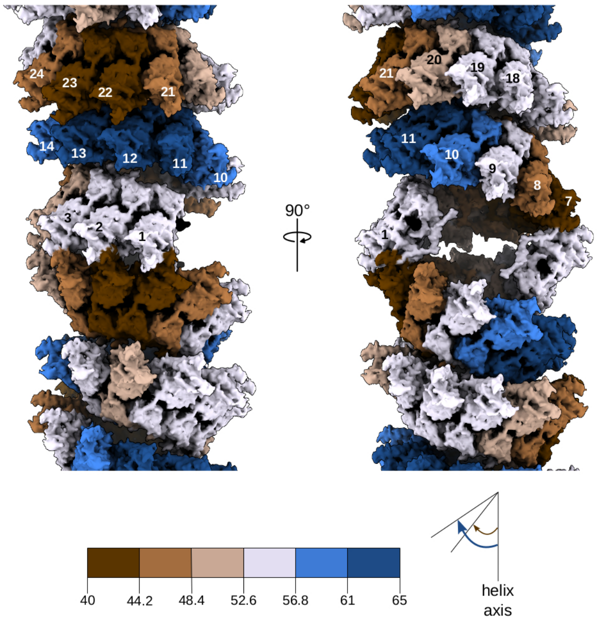


**Supplementary Figure 7. Model of a long double-headed NC**. A simulated map of long double-headed NCs, filtered to 6 Å resolution, was built from the atomic model of the double-headed NC. Protomers are coloured dependent on their axial tilt determined as described in the method section, following the colour code shown at the schematic underneath, RNA is in black. The protomer numbering follows the convention used for the non-canonical helical NC.
